# Mitochondria-Targeted OGG1 remodels Alveolar Epithelial Transitional States and Immune-Epithelial Communication after Chlorine Lung Injury

**DOI:** 10.64898/2026.10.20.760415

**Authors:** Shubham Dubey, David K Crossman, Sadis Matalon

## Abstract

Chlorine (Cl₂) inhalation causes acute epithelial and endothelial injury, pulmonary edema, inflammation, and impaired gas exchange, yet no approved therapy directly targets the mechanisms that sustain lung injury after exposure. We recently showed that oxidative mitochondrial DNA (mtDNA) damage drives chlorine-induced lung injury and that post-exposure delivery of a fusion protein containing mitochondria-targeted 8-oxoguanine DNA glycosylase-1 (mOGG1) mitigates lung injury and improves survival. However, the cell populations and transcriptional states through which mtDNA injury shapes lung damage and repair remain unclear. Here, we used single-nucleus RNA sequencing to define early cellular responses to distinct epithelial and immune cell populations. C57BL/6 mice (4/group; 2 males and 2 females per group) were exposed to air or Cl₂ (500 ppm, 30 min), returned them to room air and mOGG1 (1 mg/kg, intranasal) was administered 1 h after exposure; the mice were sacrifice and their lungs were collected at 24 h post exposure. Integrated analysis resolved 39 transcriptionally distinct epithelial, immune, endothelial, stromal, and vascular populations. Although the major cellular compartments remained broadly preserved, Cl₂ induced marked remodeling of epithelial composition and transcriptional states. Subclustering of the AT1/AT2 transitional compartment identified four Differential Transcriptomic Adaptation Phases (DTAPs): an acute stress-response state (DTAP_D), a proliferative state (DTAP_P), a Krt8⁺ damage-associated transitional state (DTAP_T), and an AT1-like resolution state (DTAP_A). Pseudotemporal analyses suggested that chlorine exposure increased heterogeneity in alveolar and airway epithelial states, with injury-associated transitional programs expanding along the inferred trajectories. Ligand-receptor analysis further predicted that Cl₂ increased the complexity of IL-1 communication among neutrophils, macrophages, and epithelial populations. mOGG1 did not globally suppress this network but instead redistributed signaling interactions, including the participation of selected DTAP populations. Together, these findings identify heterogeneous epithelial transitional states associated with Cl₂ injury and support a model in which mOGG1 promotes recovery by reshaping epithelial injury-repair programs and immune-epithelial communication.

## Introduction

Chlorine (Cl₂) is a highly reactive inhalational toxicant released during industrial accidents, transportation incidents, and occupational exposures. Acute high-dose exposure causes epithelial and endothelial injury, disruption of the alveolar-capillary barrier, pulmonary edema, inflammation, impaired gas exchange, and, in severe cases, acute respiratory distress syndrome (ARDS)^1–3.Survivors^ may also develop persistent airflow limitation and COPD-like abnormalities^4,5^ . Despite this substantial acute and chronic morbidity, no approved therapies directly target the mechanisms that sustain lung injury after chlorine exposure.

Oxidative stress is a central driver of chlorine toxicity. Chlorine reacts with water and cellular substrates to generate reactive chlorine species, including hypochlorous acid and chloramines, which oxidize proteins, lipids, and nucleic acids^3,6,7^ . Mitochondria are especially vulnerable because they are both a source and target of reactive oxygen species (ROS). Oxidative injury to mitochondrial DNA (mtDNA) can impair mitochondrial bioenergetics, promote mitochondrial ROS generation, activate regulated cell-death pathways, and lead to release of mtDNA damage-associated molecular patterns (DAMPs) that trigger innate immune signaling^8–11^ .

The base-excision repair enzyme 8-oxoguanine DNA glycosylase-1 (OGG1) initiates the repair of 8-oxoguanine (8-oxoG), a common oxidative DNA lesion^12–14^ . In our recently published report, we showed that chlorine exposure reduced lung OGG1 expression and caused prominent mtDNA damage ^15^. This was associated with mitochondrial structural abnormalities, activation of NLRP3, TLR9, and cGAS-linked inflammatory pathways, cytokine production, acute lung injury, impaired epithelial ion transport, and reduced alveolar fluid clearance. Post-exposure instillation of mitochondria-targeted OGG1 (mOGG1) initiated mtDNA damage repair, improved mitochondrial function, decreased inflammation and lung injury, restored epithelial sodium channel activity and alveolar fluid clearance, and increased survival. MitoOGG1 also attenuated longer-term COPD-like physiological changes, whereas global OGG1 deficiency worsened mortality and lung injury^16^.

These findings established oxidative mtDNA damage as a key contributor to chlorine-induced lung injury and identified mOGG1 as a promising post-exposure therapeutic strategy. However, the specific lung cell population through which mitochondrial DNA injury drives tissue damage, inflammation, and repair remain unclear. In particular, it is unknown how chlorine-induced mitochondrial dysfunction affects epithelial cell-state transitions, whether injury-associated epithelial intermediate states emerge and accumulate after exposure, and how mitoOGG1 reshapes epithelial-immune crosstalk during early repair.

In the present study, we used single-nucleus RNA sequencing to define the cellular and transcriptional responses to chlorine exposure and intranasal instillation of mitoOGG1 after chlorine exposure in the mouse lung. Building directly on our previous demonstration that mitochondrial oxidative-DNA damage is a causal contributor to chlorine lung injury^16^, we sought to identify the cellular states and intercellular signaling networks associated with the response to mitoOGG1. We identified major epithelial, immune, endothelial, stromal, and vascular populations across air-control, chlorine-exposed, and chlorine-plus-mitoOGG1 groups. Within the AT1/AT2 transitional compartment, we identified four distinct **Differential Transcriptomic Adaptation Phases (DTAPs)**: an acute stress-response state (**DTAP_D**), a proliferative state (**DTAP_P**), a Krt8-positive damage-associated transitional state (**DTAP_T**), and an AT1-like resolution-associated state (**DTAP_A**). We then used pseudotemporal analysis to place these states along an inferred alveolar injury-repair continuum and examine condition-associated changes in alveolar and airway epithelial-state organization. We also evaluated predicted IL-1-mediated communication among epithelial and immune populations. We hypothesized that, in acute chlorine lung injury, chlorine exposure would drive early remodeling of lung epithelial and immune compartments, characterized by expansion of damage-associated AT1/AT2 transitional epithelial states, activation of stress-response and proliferative transcriptional programs, and increased predicted inflammatory communication between epithelial cells and innate immune populations. Based on our prior evidence that oxidative mtDNA damage contributes to chlorine-induced lung injury and that mitoOGG1 improves physiological outcomes, we further hypothesized that a single intranasal dose of mitoOGG1 administered 1 h after Cl₂ exposure may not simply restore the lung to an air-control transcriptional state. Instead, mitoOGG1 could selectively remodel these responses and favor a more controlled injury-repair program rather than uniformly restoring the lung to its uninjured state.

## Material and methods

### Animals

Wildtype (C57BL/6) 8–12 weeks old male and female mice (∼20–25 g, body weight) were purchased from Charles River Laboratories (Wilmington, MA) and kept for at least 72h of acclimatization before using them for experiments. All experimental protocols on mice were approved by Institutional Animal Care and Use Committee of University of Alabama at Birmingham (IACUC-UAB) under APN 22366, 22606.

### Exposure to Cl2

All the exposures were done as explained previously ^16^. In short, unanesthetized male and female mice were exposed to Cl2 (500 ppm for 30 min) in Plexiglas chambers between 7 and 10 a.m.^17,18^ . According to UAB IACUC guidelines, all mice were injected with long-acting buprenorphine (1 mg/kg subcutaneously) prior to exposure to minimize anxiety and discomfort. At the end of the exposure, the mice were returned to their cages at room air and observed during the day at least every 2 h and once during the night. They were allowed access to food and water *ad libitum.* Mice were sacrificed if their body weight dropped below 30 % of their initial value.

### Generation and administration of recombinant mOGG1

A fusion protein containing OGG1 with a mitochondria targeted sequence was provided by Dr. Mark Gillespie, the University of South Alabama^19^. mOGG1 (1 mg/kg or 0.5 mg/kg in 50 μl of sterile saline) was administered in the external nares of conscious mice at 1 h post-exposure to Cl2 as described previously ^16^.

### Lung tissue collection and nuclei isolation

Whole lungs were collected at 24 h post Cl2 exposure and processed for isolation of single nuclei. Lung tissue was minced and mechanically dissociated in ice-cold nuclei isolation buffer supplemented with RNase inhibitors following manufacturer’s instruction (CG000124, Rev G). The tissue suspension was filtered through a 70um cell strainer and subjected to low-speed centrifugation to remove tissue debris. Nuclei were released by detergent-assisted lysis and washed in nuclei isolation buffer. The nuclei suspension was filtered again to remove aggregates and debris, and nuclei concentration and integrity were assessed by microscopic examination and/or automated particle counting. Nuclei were stained with a fluorescent DNA-binding dye (7AAD) and sorted by fluorescence-activated cell sorting (FACS) to enrich intact singlet nuclei and exclude debris, doublets, and damaged nuclei. 5,000 sorted nuclei were collected on ice and immediately used for library preparation.

### Single-nucleus RNA-sequencing library preparation and sequencing

Single-nucleus RNA-sequencing libraries were prepared using the 10x Genomics Chromium platform and the manufacturer’s recommended reagents and protocols (Single Cell 3’ v4 (polyA) OCM). Briefly, nuclei were partitioned into nanoliter-scale droplets containing barcoded gel beads and reverse-transcription reagents. Nucleus-derived RNA was captured using cell or nucleus specific barcodes, followed by reverse transcription, cDNA amplification, and construction of gene-expression libraries. Libraries were assessed for quality and quantified before sequencing. Libraries were sequenced using an Illumina sequencing platform with paired-end sequencing according to the recommendations for the 10x Genomics gene-expression workflow. Sequencing depth and run-specific metrics were evaluated for each library before downstream analysis.

### Processing of single-nucleus RNA-sequencing data

Sequencing data were demultiplexed and processed using the 10x Genomics Cell Ranger multi pipeline (version 9.0.1). Reads were aligned to the mouse reference genome (GRCm39-2024-A from 10X), and gene-by-nucleus expression matrices were generated for each sample. The resulting matrices were imported into R for downstream analysis using Seurat (version 5.3.1). Individual 10x Genomics datasets were imported into Seurat objects with a minimum threshold of 200 detected genes per cell and 3 cells per gene. Quality-control metrics, including total UMI counts, number of detected genes, and percent mitochondrial transcripts, were calculated for each cell. Quality-control thresholds were determined individually from the distribution of metrics within each sample to exclude low-quality cells and putative doublets. Filtered samples were merged into a single dataset for downstream analyses. Gene expression data were log-normalized, and the 2,000 most variable genes were identified using the variance-stabilizing transformation (VST) method. Data were scaled and subjected to principal component analysis (PCA). Batch correction and dataset integration were performed using Harmony through the Seurat v5 integration framework. Neighborhood graphs were constructed using the first 30 principal components, followed by Louvain clustering (resolution = 0.8) and Uniform Manifold Approximation and Projection (UMAP) for visualization.

### Nuclei-type annotation

Cell identities were assigned by reference-based annotation using the LungMAP Mouse Lung Cell Reference Atlas (v1.1). Transfer anchors were generated between the reference and query datasets using Seurat, and predicted cell-type labels were transferred to the experimental dataset. Cluster annotations were further evaluated using established marker genes and visualization of cluster-specific expression patterns. The integrated dataset resolved 39 transcriptionally distinct populations representing epithelial, stromal, vascular, and immune compartments. Cell-type identities were confirmed using feature plots, dot plots, and heatmaps of lineage-associated genes. Marker-gene heatmaps were generated using scaled expression values for representative genes across annotated populations.

### Analysis of epithelial transcriptional responses

Epithelial populations were subset from the integrated dataset for examination of treatment-associated transcriptional changes. Expression of epithelial identity, alveolar function, and injury-associated genes was visualized using heatmaps, dot plots, violin plots, and feature plots. For selected epithelial populations, expression patterns were compared among air-control, Cl₂-exposed, and Cl₂ plus mOGG1-treated groups. Differential gene expression analysis was performed using Seurat’s FindMarkers() function within these individual annotated cell populations. Results were reported using estimated log2 fold changes, p-values and adjusted P values. Because nuclei from the same mouse are not independent biological replicates, inferential comparisons were interpreted at the animal level or using pseudobulk aggregation when appropriate.

### Subclustering of the AT1/AT2 transitional compartment

AT1/AT2 cells were extracted from the integrated single-cell RNA-sequencing dataset and analyzed separately to identify damage-associated transitional alveolar progenitor (DTAP) cell states. Following normalization, dimensionality reduction, and Harmony-based integration, cells were clustered and visualized using UMAP. DTAP subpopulations were identified using the UCell package, which performs rank-based single-cell gene signature scoring. Four previously described DTAP transcriptional programs, including damage-associated (DTAP-D), alveolar-associated (DTAP-A), transitional (DTAP-T), and proliferative (DTAP-P) states, were evaluated using published DTAP gene signatures. DTAP-D, DTAP-A, DTAP-T, and DTAP-P gene signatures were derived from previously published descriptions of damage-associated transitional alveolar progenitor cell states. UCell scores were calculated independently for each signature, and cells were assigned to the DTAP state corresponding to the highest-confidence signature score. Cells with ambiguous assignments were excluded from DTAP-specific analyses. DTAP annotations were incorporated into the Seurat object and used for downstream analyses, including visualization of DTAP distributions across treatment groups, identification of DTAP-associated marker genes, and differential gene expression analyses between Air Control, Cl₂-exposed, and Cl₂+OGG1 treatment groups within each DTAP state.

### Pseudotemporal analysis of DTAP states

Pseudotime analysis was performed on confidently classified DTAP populations using Monocle3. Cells assigned to DTAP-A, DTAP-D, DTAP-P, or DTAP-T states by UCell were extracted from the AT1/AT2 subset, while cells with ambiguous DTAP assignments were excluded from trajectory analyses. The DTAP subset was converted from a Seurat object to a Monocle3 cell data set, and experimental group metadata were retained for downstream analyses. Cells were preprocessed using principal component analysis (30 dimensions), and the UMAP embedding generated in Seurat was transferred to Monocle3 for trajectory inference. Cells were clustered within Monocle3, and a principal graph representing the developmental trajectory was learned using a joint trajectory framework across all experimental groups. To establish biological directionality, DTAP-D cells were designated as the root population for pseudotime ordering. Pseudotime values were subsequently calculated for all cells relative to the DTAP-D state. Trajectory structure and pseudotime distributions were visualized across Air Control, Cl₂-exposed, and Cl₂+OGG1 treatment groups. To identify genes associated with progression along the inferred trajectory, Monocle3 graph-based testing was performed using the principal graph, and genes exhibiting significant pseudotime-dependent expression patterns were identified based on false-discovery-rate-adjusted significance thresholds. Expression dynamics of selected genes were visualized as smoothed trends across pseudotime.

### Separate alveolar and airway trajectory analyses

Monocle3 was used to reconstruct epithelial lineage trajectories within airway and alveolar compartments. Airway analyses were performed on Basal, Secretory, Ciliated, and Deuterosomal cells, whereas alveolar analyses included AT2, AT1/AT2, AT1, DTAP-A, DTAP-D, DTAP-P, and DTAP-T populations. Subsetted Seurat objects were converted to Monocle3 cell data sets, and the Seurat-derived UMAP embeddings were transferred to Monocle3 for trajectory inference. Following preprocessing, clustering, and principal graph learning, cells were ordered along pseudotime using Basal cells as the root population for airway trajectories and AT2 cells as the root population for alveolar trajectories. Joint trajectories were constructed using cells from all experimental groups. Trajectory structure, pseudotime distributions, and treatment-associated effects were evaluated across Air Control, Cl₂-exposed, and Cl₂ plus OGG1 groups. Genes exhibiting significant pseudotime-dependent expression patterns were identified using Monocle3 graph-based testing and visualized across pseudotime.

### Cell-cell communication analysis

Cell-cell communication analysis was performed using the **CellChat** package on a subset of epithelial and immune cell populations identified from the integrated single-cell RNA-sequencing dataset. Epithelial populations included AT1, AT2, AT1/AT2, and Secretory cells, while immune populations included alveolar macrophages (AM), interstitial macrophages (IM), inflammatory monocytes (iMON), and neutrophils. Analyses were conducted separately for Air Control, Cl₂-Exposed, and Cl₂+OGG1 treatment groups. For each treatment group, CellChat objects were constructed from the Seurat object using cell-type annotations as communication nodes and the CellChat mouse ligand-receptor database as the interaction reference. Signaling genes and overrepresented ligand-receptor interactions were identified, and intercellular communication probabilities were inferred using a truncated mean model. Interactions involving cell populations containing fewer than 10 cells were excluded. Communication networks were subsequently aggregated at both the ligand-receptor and signaling pathway levels, and pathway-specific network centrality scores were calculated to identify major signaling sources and targets. To compare communication networks across experimental groups, CellChat objects were merged and analyzed using CellChat’s comparative framework. Differences in the number and strength of predicted interactions, signaling pathway information flow, ligand-receptor activity, and signaling roles of individual cell populations were assessed between Air Control, Cl₂-Exposed, and Cl₂+OGG1 groups. Network similarity analyses were additionally performed to evaluate treatment-associated changes in overall communication architecture.

### Statistical analysis

For animal-level analyses, the mouse was considered the experimental unit. Data was assessed for distribution and analyzed using statistical tests appropriate for the experimental design. Comparisons among air-control, Cl₂-exposed, and Cl₂ plus mOGG1-treated groups were performed using one-way analysis of variance followed by an appropriate multiple-comparison test, or by a nonparametric equivalent when normality assumptions were not met. Sex was balanced across groups and was included as a biological variable where the sample size and statistical model permitted. For single-nucleus RNA-sequencing analyses, differential expression and cell-type abundance comparisons were performed using biological replicates or pseudobulk aggregation to minimize pseudoreplication. Multiple-testing correction was performed using the Benjamini-Hochberg method. Adjusted P values less than 0.05 were considered statistically significant unless otherwise stated.

## Results

### Single-nuclei RNA sequencing resolves diverse cellular populations in the mouse lung

Chlorine exposure was performed at 500 ppm for 30 min, and mOGG1 was intranasally instilled at a dose of 1 µg/g body weight 1 h post-exposure. Lungs were collected 24 h after exposure to chlorine and single nuclei were extracted from four mice (two male and two female) from each group.

The experimental and computational workflow is illustrated in **(Figure 1A)**. Nuclei were isolated from whole-lung tissues, sorted by fluorescence-activated cell sorting (FACS), and 5000 nuclei per mouse were processed for library construction using the 10x Genomics platform (Single Cell 3’ v4 (polyA) OCM). Following sequencing on the Illumina platform with 40 million reads/5000 nuclei and gene-expression quantification, nuclei were grouped by transcriptional similarity and annotated according to established marker-gene expression patterns.

**Figure 1.**
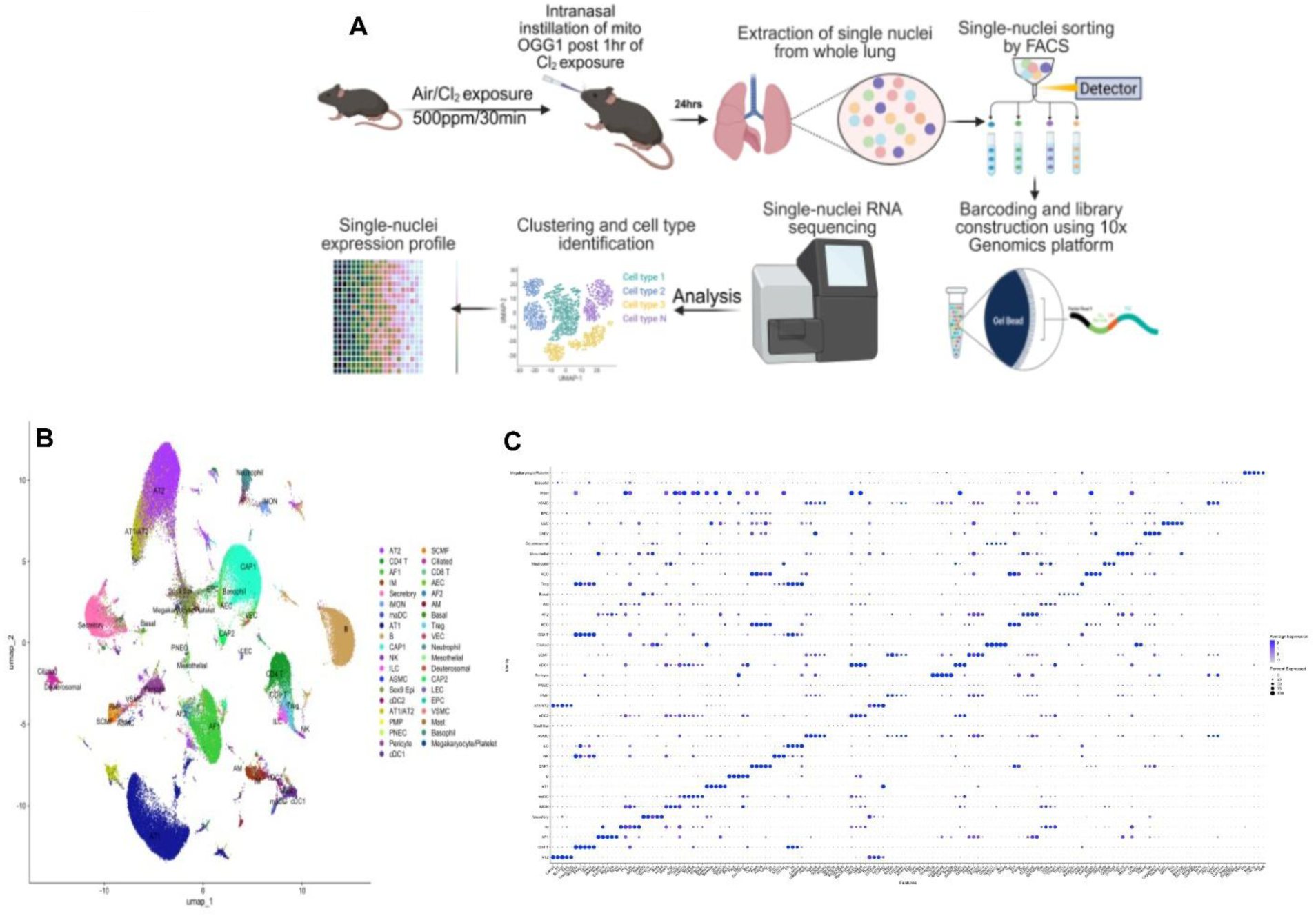
Experimental workflow and single-nuclei transcriptomic profiling of the mouse lung. **(A)** Schematic of the experimental and single-nuclei RNA-sequencing workflow. Mice were exposed to air or chlorine at 500ppm for 30 min, followed by intranasal administration of mitochondria-targeted OGG1 (mOGG1,1 µg/g body weight) 1 h post chlorine exposure. Whole lungs were collected 24 h after exposure, and nuclei were isolated from lung tissue. Nuclei were sorted by fluorescence-activated cell sorting, barcoded, and processed for library preparation using the 10x Genomics platform (Single Cell 3’ v4 (polyA) OCM). Single-nuclei RNA sequencing was followed by generation of gene-expression profiles, clustering, cell-type identification, and downstream transcriptomic analysis. **(B)** UMAP visualization of the single-nuclei RNA-sequencing dataset from whole-lung tissue. Each point represents an individual nucleus and is colored according to its assigned cell type. The dataset resolved diverse epithelial, stromal, vascular, and immune populations, including alveolar type 1 and type 2 epithelial cells (AT1 and AT2), ciliated, basal, secretory, and pulmonary neuroendocrine cells; fibroblast populations (AF1 and AF2); endothelial, lymphatic endothelial, pericyte, and vascular smooth muscle populations; and immune populations, including alveolar and interstitial macrophages, monocytes, dendritic cells, neutrophils, natural killer cells, innate lymphoid cells, B cells, CD4-positive T cells, CD8-positive T cells, regulatory T cells, mast cells, basophils, and megakaryocyte/platelet-associated cells. Cell identities were assigned based on established lineage-associated transcriptional markers. **(C)** Dot plot showing scaled expression of representative marker genes across individual nuclei grouped by annotated cell type. Columns are separated by marker genes/features, and rows represent genes used to define cellular identity. Dot size represent the percentage of cells expressing the gene . The distinct marker-gene patterns across the clusters support the cell-type assignments shown in **panel B**.

The UMAP projection demonstrated substantial transcriptional heterogeneity with 39 different cell clusters across the lung and resolved a broad range of epithelial, stromal, vascular, and immune populations **(Figure 1B)**. Epithelial clusters included AT1 and AT2 cells, as well as ciliated, basal, secretory, pulmonary neuroendocrine, and deuterosomal populations. Stromal and vascular compartments included AF1 and AF2 fibroblast populations, capillary and other endothelial populations, lymphatic endothelial cells, pericytes, vascular smooth muscle cells, and mesothelial cells. The dataset also contained multiple immune-cell populations, including alveolar and interstitial macrophages, monocytes, dendritic cells, neutrophils, natural killer cells, innate lymphoid cells, B cells, CD4-positive T cells, CD8-positive T cells, and regulatory T cells. Smaller populations, including mast cells, basophils, and megakaryocyte/platelet-associated cells, were also identified. The separation of these populations in UMAP space indicates that the sequencing approach captured the major cellular compartments of the mouse lung.

### Marker-gene expression supports cell-type annotation

The heatmap demonstrated distinct expression patterns across the annotated populations **(Figure 1C)**.AT1 and AT2 epithelial cells displayed separate transcriptional signatures, while ciliated, basal, secretory, and pulmonary neuroendocrine populations exhibited their own characteristic marker profiles. Similarly, fibroblast, endothelial, pericyte, smooth muscle, and mesothelial clusters showed compartment-specific gene-expression patterns.

Immune populations were also distinguished by lineage-associated transcriptional programs. Macrophage, monocyte, dendritic-cell, neutrophil, lymphoid, mast-cell, basophil, and megakaryocyte/platelet-associated clusters each display distinguishable expression patterns. Overall, the concordance between the UMAP organization and marker-gene expression supports the identification of the major lung cell types and establishes a cellular framework for examining cell-type-specific responses to chlorine-induced lung injury and the potential protective effects of mOGG1.

### Single-nuclei RNA-seq identifies chlorine-induced changes in lung cellular composition and epithelial transcriptional states that are restored by mito-OGG1

To define the early lung response to chlorine inhalation and determine the effect of mitochondria-targeted OGG1, we analyzed lungs collected 24h after exposure from air-control, Cl_2_-exposed, and Cl_2_-exposed mice treated with mito-OGG1. Integrated single-nuclei RNA-seq analysis identified multiple epithelial, immune, endothelial, and stromal populations across all groups (**Figure 2A**).

**Figure 2.**
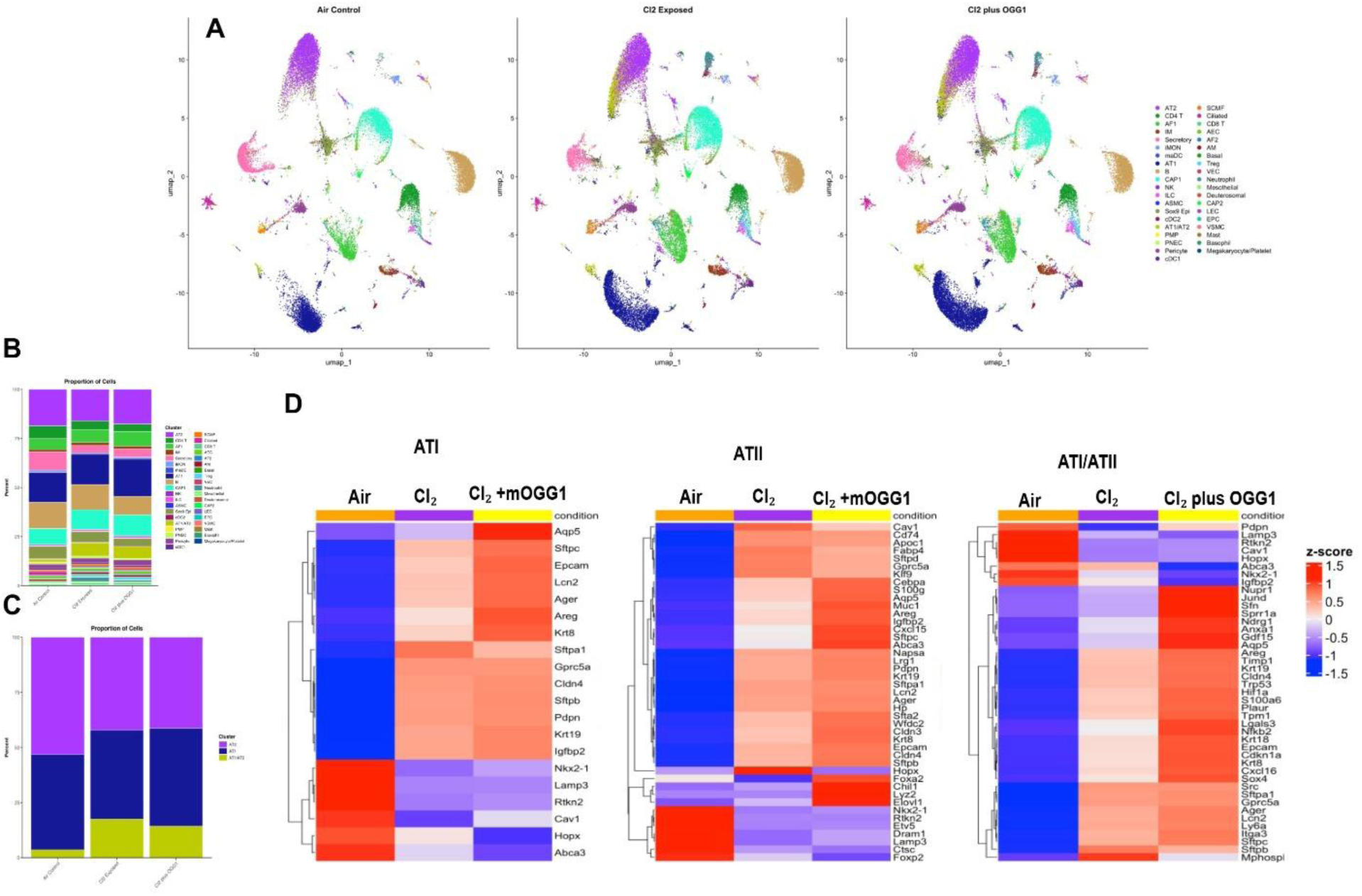
Single-nucleus RNA-seq analyses reveal chlorine-induced epithelial remodeling and modulation by mOGG1. **(A)** UMAP visualization of integrated single-nucleus RNA-sequencing data from lungs collected 24 h after treatment. C57BL/6 mice were exposed to room air or chlorine (500 ppm for 30 min), and the treatment group received mitochondria-targeted OGG1 intranasally (1 µg/g body weight) 1 h after chlorine exposure. Nuclei are colored by annotated cell identity and shown separately for air control, Cl2-exposed, and Cl2 plus mOGG1 groups. Major epithelial, immune, endothelial, and stromal populations were identified across all conditions. **(B)** Stacked bar plots showing the relative abundance of annotated cell populations in each experimental group. Chlorine exposure altered the proportional representation of multiple lung cell populations, whereas mOGG1 treatment partially shifted these changes toward the air-control distribution. **(C)** Stacked bar plots show the percentages of alveolar type 1 (AT1), alveolar type 2 (AT2), and transitional AT1/AT2 cells among total alveolar epithelial cells in the air-control, Cl₂-exposed, and Cl₂ plus mOGG1 groups. Each color denotes a distinct cell population, and each bar sums to 100%. **(D)**Heatmaps showing z-score-normalized average expression of representative genes in selected alveolar epithelial populations from the single-nucleus RNA-seq dataset. In AT1/AT2-related epithelial populations, chlorine exposure increased expression of injury- and transitional-state-associated genes, including **Clip4, Ager, Sftpb, Krt8, Krt19,** and **Krt18**, and mOGG1 shifted these chlorine-induced changes toward the air-control pattern. We also observed changes in epithelial identity and stress-response genes, consistent with chlorine-induced epithelial remodeling.

The overall topology of the UMAP plots was similar across the three treatment groups, indicating that chlorine exposure and subsequent mOGG1 administration did not substantially alter the major lung cell populations or produce a gross reorganization of the global cellular landscape. Nevertheless, differences in the density and distribution of nuclei within individual clusters were apparent between conditions, suggesting that chlorine exposure may alter the relative representation and/or transcriptional state of specific lung cell populations. The mOGG1-treated group generally retained the overall cellular architecture observed in the control and chlorine-exposed datasets, indicating that its protective effects are more likely reflected in cell-state-specific transcriptional responses and shifts in population representation rather than in the emergence or loss of major cell types.

The relative abundance of the major cell populations was generally similar across the three groups, although several differences were apparent (**Figure 2B**). Compared with air controls, chlorine exposure was associated with a lower proportion of secretory, AT2, CD4 T and Sox9⁺ epithelial cells and a modest increase in the relative representation of AT1/AT2 intermediate, CAP1 cells. The mOGG1-treated group retained the overall cellular composition observed after chlorine exposure, while partially attenuating some chlorine-associated changes and increasing the relative representation of AT1, AT2, and AF1 population. Within the alveolar epithelial compartment, AT2 cells accounted for approximately 53% (**Figure 2C**) of cells in air-control lungs but decreased to approximately 41% after chlorine exposure. In parallel, the AT1/AT2 intermediate population increased from approximately 4% in air controls to approximately 18% following chlorine exposure, consistent with a shift toward an intermediate alveolar epithelial state. AT1 cells represented approximately 43% of the alveolar epithelial compartment in controls and approximately 41% after chlorine exposure.

mOGG1 treatment partially attenuated the chlorine-associated expansion of the AT1/AT2 intermediate population, which decreased to approximately 14%, while the AT1 population increased to approximately 45%. The proportion of AT2 cells remained approximately 41% after mOGG1 treatment, similar to the chlorine-exposed group and lower than in air controls. Thus, at this early time point and with this treatment regimen, mOGG1 treatment was associated with a relative redistribution of alveolar epithelial populations, characterized primarily by a reduction in the AT1/AT2 intermediate compartment and a modest increase in AT1 representation, rather than complete restoration of the control AT2 proportion.

To further examine condition-associated molecular changes, we generated gene-expression heatmaps for selected epithelial populations **(Figure 2D)**. The heatmaps showed distinct expression patterns across the air control, chlorine-exposed, and chlorine plus mOGG1 groups. The analyzed populations expressed epithelial and alveolar-associated genes, including *Epcam*, *Krt8*, *Krt19*, *Sftpc*, *Sftpb*, *Sftpa1*, *Abca3*, *Ager*, *Aqp5*, *Pdpn*, *Cav1*, and *Hopx*, supporting their epithelial and alveolar identities.

Within the AT1 epithelial population, chlorine exposure increased expression of several epithelial injury- and stress-associated genes, including Lcn2, Areg, Krt8, and Krt19, along with changes in alveolar marker expression such as *Sftpc*, *Sftpb*, and *Ager* **(Figure 2D)**. At the same time, canonical AT1-associated genes, including *Cav1*, *Hopx*, and *Abca3*, showed altered expression relative to air controls. The chlorine plus mOGG1 group showed a modified expression pattern relative to chlorine alone, with several genes displaying expression levels that were intermediate between, or closer to, those observed in air controls. The ATII epithelial population showed a similar pattern **(Figure 2D)**. Chlorine exposure altered genes involved in epithelial identity, alveolar function, and injury responses, including *Cd74*, *Fabp4*, *Sftpc*, *Krt19*, *Lcn2*, *Ager*, and *Epcam*. Expression patterns in the mOGG1-treated group differed from those in the chlorine-only group, consistent with a treatment-associated shift in the epithelial transcriptional response.

The ATI/ATII transitional epithelial population also demonstrated condition-dependent changes in gene expression **(Figure 2D)**. Genes associated with alveolar epithelial identity and differentiation, including *Pdpn*, *Cav1*, *Hopx*, *Abca3*, *Ager*, and *Sftpc*, varied across the treatment groups. Chlorine exposure altered the expression of several epithelial marker genes, whereas mOGG1 treatment further shifted this transcriptional program. Collectively, these findings indicate that chlorine inhalation induces molecular remodeling of AT1, AT2, and transitional alveolar epithelial populations. mOGG1 modifies these responses, but its effect is not represented by a simple return of all genes to control levels. Instead, the data supports treatment-associated reprogramming of epithelial cell states, potentially involving pathways related to epithelial stress, injury, differentiation, and repair.

Taken together, the UMAP and heatmap analyses indicate that chlorine exposure induces transcriptional changes in distinct lung epithelial populations without disrupting the overall cellular composition of the lung. The altered expression of epithelial injury, differentiation, and alveolar function-associated genes, together with the modified profiles observed after mOGG1 treatment, supports a potential protective or restorative effect of mOGG1 on chlorine-associated epithelial injury. Formal determination of treatment effects will require quantitative comparisons of cell abundance and differential gene expression across biological replicates.

### Subclustering of the AT1/AT2 Transitional Compartment identifies distinct Differential Transcriptomic Adaptation Phases following Chlorine Injury

To further define the epithelial response to chlorine injury, we subclustered the AT1/AT2 transitional compartment, a population that in recent single-cell atlases of injured lung epithelium has been shown to contain damage-associated intermediate states co-expressing canonical alveolar markers together with stress, proliferative, and differentiation programs. This analysis resolved four reproducible transcriptional states, which we designate Differential Transcriptomic Adaptation Phases (DTAPs), showed mixed features. Notably, DTAP states were markedly enriched in the Cl₂ exposed and Cl₂ plus mOGG1 groups and were only rarely detected in air-control lungs (**Figure 3A**).

**Figure 3.**
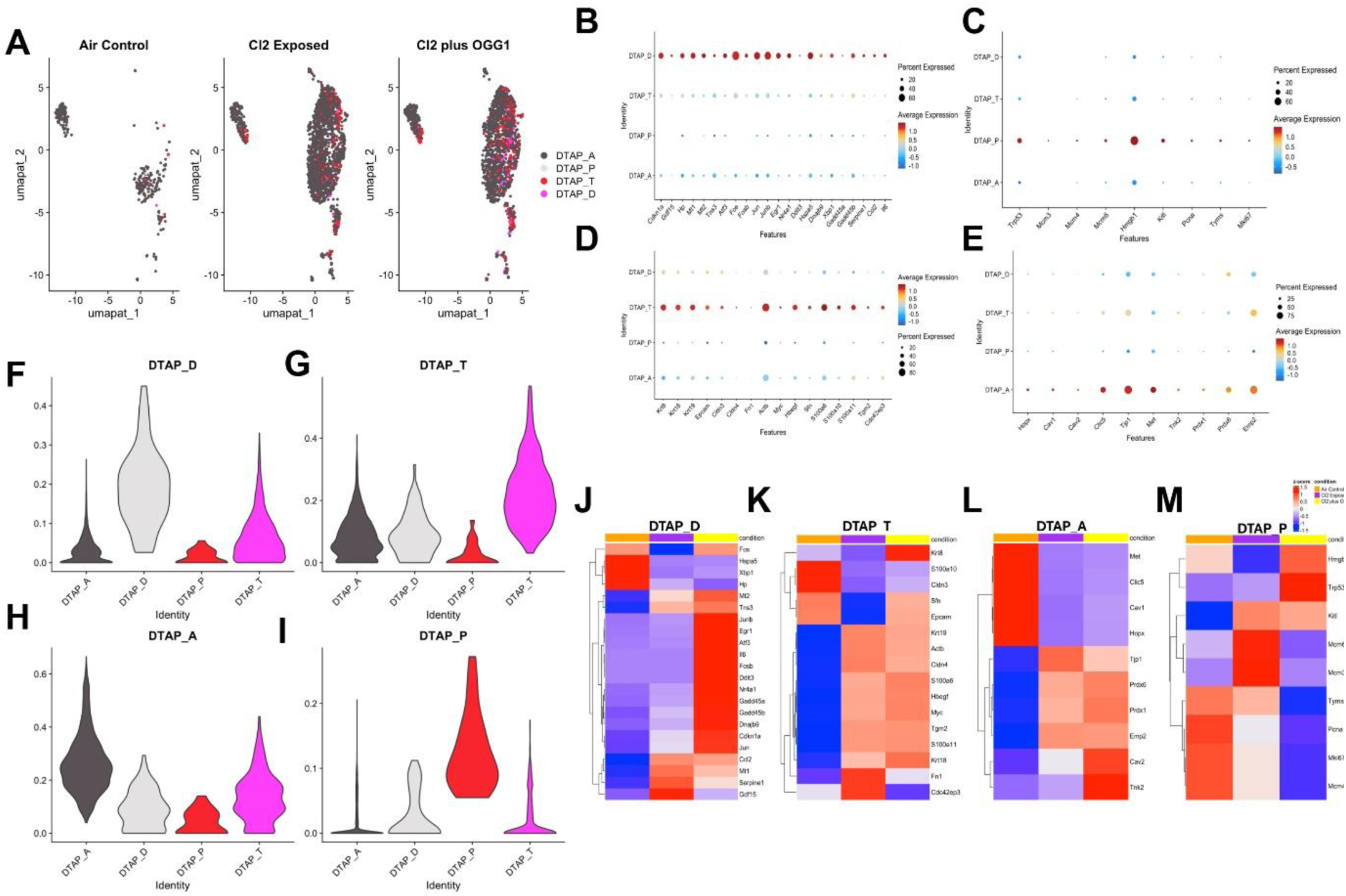
Subclustering of the AT1/AT2 transitional compartment identifies four Differential Transcriptomic Adaptation Phases (DTAPs) after chlorine injury. Mice were exposed to 500 ppm chlorine for 30 min, treated with intranasal mitochondria-targeted OGG1 (1 µg/g body weight) 1 h after exposure, and lungs were collected 24 h later. Each group contained 4 mice (2 males, 2 females). **(A)** UMAP visualization of subclustered alveolar epithelial transitional cells from air control, Cl_2_-exposed, and Cl_2_ plus mOGG1-treated lungs. Cells are colored by DTAP identity. Based on marker expression and module scoring, the four resolved states correspond to: DTAP_D, an acute activation/stress-response phase; DTAP_P, a proliferative/transient progenitor phase; DTAP_T, a damage-associated transitional Krt8^+^/PATS/DATP-like phase; and DTAP_A, a resolution/terminal differentiation phase toward AT1 identity. An additional Ambiguous cluster contained cells with mixed or intermediate transcriptional features. **(B)** Dot plot of representative acute stress/injury-response genes identifying the DTAP_D, (**C**) Dot plot of representative cell-cycle/proliferation-associated genes identifying the DTAP_P state. **(D)** Dot plot of representative damage-associated transitional genes, including Krt8/PATS-like markers, identifying the DTAP_T state. (**E**) Dot plot of representative AT1 differentiation/resolution-associated genes identifying the DTAP_A state. (**F–I**) Violin plots showing module scores for each DTAP program across the indicated cell identities, confirming selective enrichment of the cognate transcriptional signature in each subcluster. **(J-M)** Condition-dependent gene-expression patterns across DTAP cell states. Heatmaps show the scaled average expression (row z-score) of selected marker genes in DTAP_D **(J**), DTAP_T(K), DTAP_A (**L**), and DTAP_P (**M**) cells from the air-control, Cl₂-exposed, and Cl₂ plus mOGG1 groups. Columns represent experimental conditions, indicated by the top annotation (orange, air control; purple, Cl₂ exposed; yellow, Cl₂ plus OGG1), and genes are hierarchically clustered by expression pattern. Red indicates higher and blue indicates lower relative expression (z-score range, -1.5 to 1.5).

These four DTAPs appear to represent a biologically ordered spectrum of epithelial adaptation after injury. DTAP_D corresponded to an early acute activation phase, characterized by strong expression of immediate-early and stress-response genes including *Atf3, Fos, Fosb, Jun, Junb, Egr1, Nr4a1, Ddit3, Hspa5, Dnajb9, Xbp1, Gadd45a, and Gadd45b* **(Figure 3B)**. This profile is consistent with activation of stress-response pathways, including unfolded-protein and endoplasmic-reticulum stress, and identifies DTAP_D as an acute injury-associated epithelial state ^18,20,21^. A second cluster, DTAP_P, represented a proliferative/transient progenitor phase. This state was defined by enrichment of cell-cycle and DNA replication genes, including Mcm3, Mcm4, Mcm6, Pcna, Tyms, and Mki67 **(Figure 3C)**. This expression pattern is consistent with a proliferative epithelial state with transient progenitor-like features^22–25^ . However, the marker analysis alone does not establish whether DTAP_P cells are more abundant after Cl₂ exposure or after mito-OGG1 treatment. The third state, DTAP_T, corresponded closely to the damage-associated transitional Krt8^+^ population previously described as PATS/DATP-like intermediates in injured lung and fibrotic lungs ^24–28:^ DTAP_T cells showed high expression of Krt8, Krt18, Krt19, Cldn3, Cldn4, Sfn, Hbegf, Cd44, Tgm2, and S100a6/S100a10/S100a11 (**Figure 3D**), consistent with a stable transitional epithelial state associated with injury remodeling. This population also showed features compatible with stress-associated remodeling and partial senescence-like signaling^26,28^ , supporting the idea that DTAP_T may represent a key intermediate or potential bottleneck in alveolar repair.

Finally, DTAP_A was enriched for genes associated with AT1 differentiation and epithelial resolution, including Hopx, Cav1, Cav2, Clic5, Tjp1, Emp2, Prdx1, and Prdx6 (**Figure 3E**). This expression pattern is consistent with progression toward restoration of a mature AT1-like identity, representing a late resolution phase in which transitional and proliferative programs are attenuated, and alveolar epithelial architecture is re-established.

The violin plots confirmed that each subcluster was selectively enriched for its corresponding DTAP signature, supporting the robustness of this classification (**Figure 3F-I**). Collectively, these findings define four transcriptionally distinct states within the AT1/AT2 transitional epithelial compartment: an acute stress-responsive state (DTAP_D), a proliferative state (DTAP_P), a damage-associated transitional state (DTAP_T), and an AT1-like differentiated state (DTAP_A).

### mOGG1 modifies chlorine induced transcriptional programs across transitional epithelial cell states

The condition-specific heatmaps further demonstrated that the four DTAP states were transcriptionally modified across treatment groups (**Figure 3J**). In DTAP_D, stress-response genes such as *Atf3*, *Fosb*, *Jun*, *Egr1*, *Ddit3*, *Gadd45a*, *Gadd45b*, *Dnajb9*, and *Cdkn1a* were most prominent in the Cl₂ plus mOGG1 group, whereas *Ccl2*, *Mt1*, *Serpine1*, and *Gdf15* were relatively more pronounced after chlorine exposure alone. This pattern suggests that mOGG1 altered the composition of the acute stress-response program rather than uniformly suppressing it.

The DTAP_T heatmap showed condition-dependent changes in transitional epithelial markers. Several genes, including *Krt19*, *Actb*, *Cldn4*, *S100a6*, *Hbegf*, *Myc*, *Tgm2*, and *S100a11*, were increased in the Cl₂ exposed group and remained elevated or were further modified following mOGG1 treatment (**Figure 3K**). In contrast, *Krt8*, *S100a10*, and *Cldn3* showed a different pattern, with lower expression after chlorine exposure and partial recovery or modification in the Cl₂ plus mOGG1 group. These findings indicate that mito-OGG1 reshaped, but did not completely normalize, the injury-associated transitional epithelial program.

Within DTAP_A, canonical AT1-associated genes, including *Met*, *Clic5*, *Cav1*, and *Hopx*, were most highly expressed in air-control cells and reduced following chlorine exposure (**Figure 3L**). mOGG1 did not uniformly restore these genes to control levels. However, other genes associated with epithelial integrity and redox regulation, including *Tjp1*, *Prdx1*, *Prdx6*, and *Emp2*, were increased after chlorine exposure and remained elevated or were partially modified in the miOGG1 group. *Cav2* and *Tnk2* were relatively more prominent after mOGG1 treatment, suggesting selective remodeling of the AT1-associated transcriptional program.

The DTAP_P heatmap showed the clearest association with chlorine-induced proliferation. Cell-cycle and DNA-replication genes, including *Mcm3*, *Mcm4*, *Mcm6*, *Pcna*, *Tyms*, and *Mki67*, were generally increased in the Cl₂-exposed group, whereas their expression was lower in the Cl₂+mOGG1 group (**Figure 3M**). This pattern suggests that mito-OGG1 attenuates the chlorine-associated proliferative program within the DTAP_P state. Together, these heatmaps show that chlorine exposure induces distinct transcriptional changes across acute stress-responses, transitional, AT1-like, and proliferative epithelial states. mOGG1 modified each of these programs, including attenuation of the chlorine-associated proliferative signature and selective restoration or reorganization of stress, transitional, and AT1-associated gene expression.

### Pseudotemporal analysis supports progression of AT1/AT2 transition cells through sequential DTAP states

To determine whether AT1/AT2 transition cells undergo an ordered progression through distinct damage-associated transitional alveolar progenitor states, we performed pseudotemporal trajectory analysis using DTAP_D cells as the trajectory origin. DTAP_D was defined as the acute activation/stress-response state, followed by DTAP_T, the damage-associated transitional Krt8⁺/PATS/DATP-like state, DTAP_P, the proliferative or transient progenitor state, and DTAP_A, the resolution/terminal differentiation state associated with progression toward AT1 identity. In air-control lungs, DTAP_D cells localized to the earliest pseudo time position, consistent with their designation as the initiating stress-response state (**Figure 4A**). DTAP_T cells occupied an early-to-intermediate pseudotime position, while DTAP_P cells were distributed at later pseudotime values. DTAP_A cells showed the broadest and generally latest pseudotime distribution, consistent with their position as a terminal or resolution-associated state. Overall, the pseudotime distribution in control lungs was compatible with an ordered transition from DTAP_D through DTAP_T toward downstream DTAP_P and DTAP_A states. Chlorine exposure altered the distribution of AT1/AT2 transition states along pseudotime (**Figure 4B**). DTAP_D cells remained concentrated near the trajectory origin, whereas DTAP_T cells displayed a broader pseudotime distribution extending into intermediate and later regions. This pattern is consistent with accumulation and expansion of the damage-associated transitional state following chlorine-induced injury. DTAP_A cells also extended across late pseudotime values, including a subset of cells at particularly late positions, suggesting increased representation of cells progressing toward a resolution or terminal differentiation state. DTAP_P cells remained primarily in the early-to-intermediate pseudotime range, although some cells occupied later positions.

**Figure 4.**
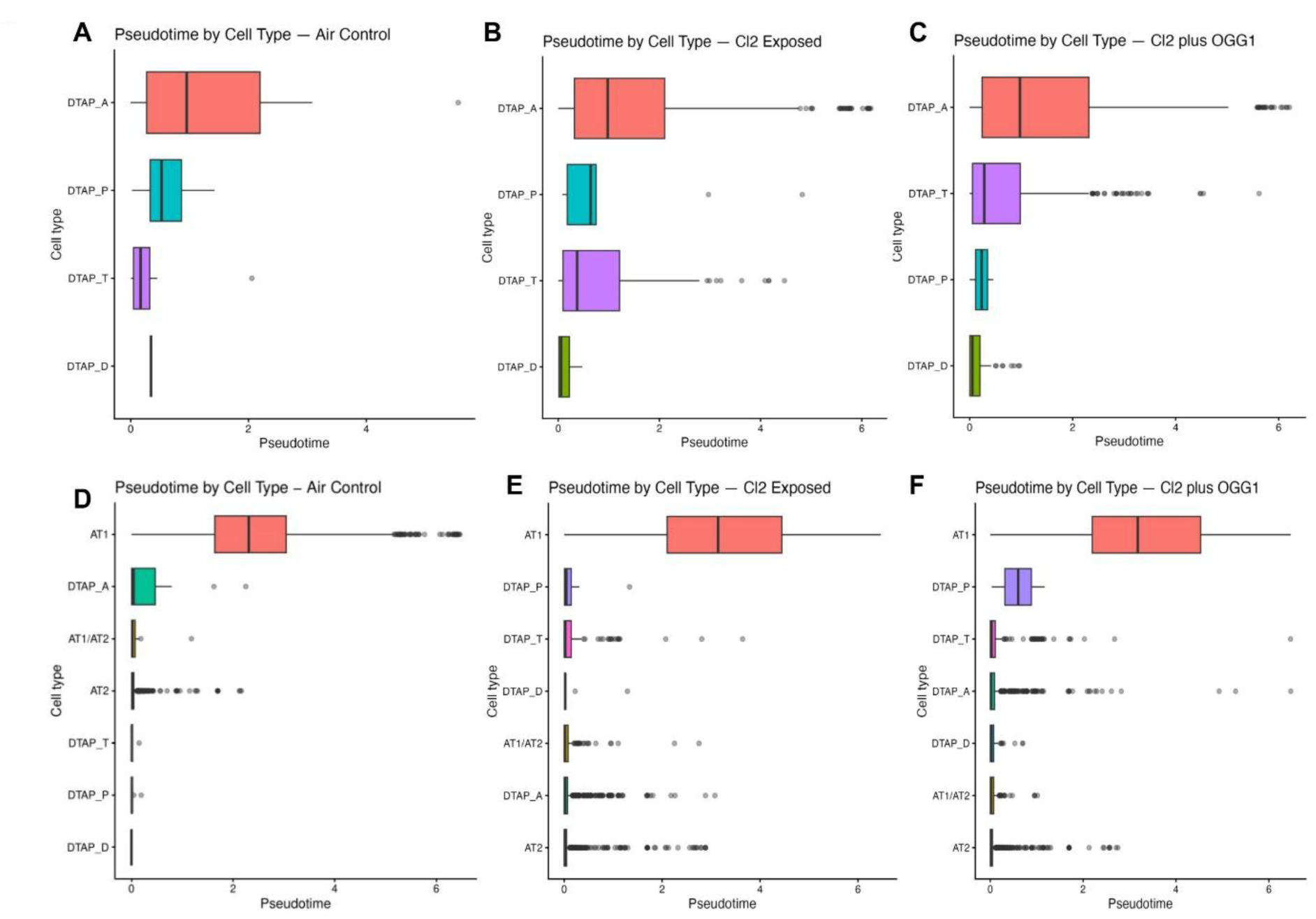
Pseudotemporal progression of AT1/AT2 transition cells through DTAP phases after chlorine exposure and mOGG1 treatment. (A-C) Pseudotemporal trajectory analysis was performed on AT1/AT2 transition cells to evaluate the progression of damage-associated transitional alveolar progenitor (DTAP) states. Pseudotime was initialized using DTAP_D as the origin. DTAP_D represents the acute activation/stress-response phase; DTAP_T represents the damage-associated transitional Krt8⁺/PATS/DATP-like phase; DTAP_P represents the proliferative or transient progenitor phase; and DTAP_A represents the resolution/terminal differentiation phase associated with progression toward AT1 identity. Boxplots show the distribution of pseudotime values for each DTAP phase in air-control, Cl₂-exposed, and Cl₂ plus mOGG1-treated lungs. (**A**) In the air-control group, DTAP_D cells occupied the earliest pseudotime position, followed by DTAP_T and DTAP_P cells, whereas DTAP_A cells were distributed across later pseudotime values. **(B)** Chlorine exposure broadened the pseudotime distribution of DTAP_T cells and increased the representation of late DTAP_A cells. **(C)** Following mOGG1 treatment, the DTAP phases retained an ordered pseudotemporal arrangement, with DTAP_D and DTAP_P cells remaining near the early pseudotime region, DTAP_T cells occupying an intermediate position, and DTAP_A cells extending toward the late pseudotime region. These findings are consistent with progression from DTAP_D through DTAP_T toward DTAP_P and/or DTAP_A, with DTAP_A and DTAP_P representing resolution/differentiation and progenitor-associated endpoints, respectively. **(D-F) Pseudotime distribution of alveolar epithelial cell states across treatment groups.** Box-and-whisker plots show pseudotime values stratified by cell type/state in air control mi **(D),** chlorine-exposed mice **(E),** and chlorine-exposed mice treated with mOGG1 (**F**). Cell states include AT1, AT2, AT1/AT2 transitional cells, and distinct damage-associated transient alveolar progenitor subclusters: DTAP_A, DTAP_D, DTAP_P, and DTAP_T. Boxes represent the interquartile range with the median indicated by the central line; whiskers denote the non-outlier range, and individual points indicate cells with more extreme pseudotime values. Chlorine exposure is associated with expansion and redistribution of transitional/DTAP populations along the pseudotime trajectory, whereas Cl₂ + mOGG1 alters the distribution of DTAP states relative to chlorine exposure alone.

In the Cl₂ plus mOGG1 group, DTAP_D cells remained near the pseudotime origin, while DTAP_T and DTAP_P cells were concentrated predominantly within early-to-intermediate pseudotime regions (**Figure 4C**). DTAP_A cells continued to occupy the later pseudotime region, preserving the expected position of the resolution/terminal differentiation state. Compared with the Cl₂-exposed group, mito-OGG1 treatment was associated with a more organized distribution of the DTAP phases and reduced broadening of the transitional/progenitor-associated populations. These findings suggest that mOGG1 does not eliminate the DTAP response but may modulate its progression and promote more orderly movement from the acute stress state through the transitional phase toward progenitor-associated or resolution-associated endpoints. Collectively, the pseudotemporal analysis supports a model in which AT1/AT2 transition cells progress from DTAP_D → DTAP_T → DTAP_P and/or DTAP_A. The DTAP_P and DTAP_A states likely represent distinct downstream outcomes, corresponding to a transient proliferative/progenitor program and a resolution/terminal differentiation program toward AT1 identity, respectively. Chlorine exposure was associated with altered and broadened progression through these states, particularly within the DTAP_T compartment, whereas mOGG1 treatment partially restored a more ordered pseudotemporal pattern. Because pseudotime represents an inferred temporal ordering rather than a direct time-course measurement, these findings should be interpreted as evidence of transcriptional state progression rather than definitive lineage tracing.

### Pseudotemporal analysis positions DTAP states along an alveolar epithelial injury-repair continuum

To examine epithelial injury-to-repair dynamics at single nucleus resolution, we performed pseudotemporal analyses and an alveolar trajectory was generated using AT2, AT1, AT1/AT2 transitional, and DTAP populations, with quiescent AT2 cells from air-control lungs used to define the trajectory origin. In air-control lungs, AT2 cells and the DTAP populations were concentrated near the early pseudotime region, whereas AT1 cells occupied later pseudotime values (**Figure 4D**). This organization was consistent with a trajectory beginning in an AT2-like state and extending toward a more differentiated AT1-like state. AT1 cells showed a broad distribution across later pseudotime values, while AT1/AT2 transitional cells were primarily localized near the early portion of the trajectory.

Following chlorine exposure, the pseudotemporal organization of the alveolar lineage was altered (**Figure 4E**). AT1 cells displayed a broader distribution extending across intermediate and late pseudotime values, indicating increased heterogeneity in the AT1-associated transcriptional state. Several DTAP populations also showed broader distributions and late-pseudotime outliers compared with air controls. These changes were consistent with chlorine-associated disruption of the normal alveolar state landscape and the emergence of heterogeneous injury-associated transitional states. The Cl₂ plus mOGG1 group showed a distinct distribution of alveolar states (**Figure 4F**). AT1 cells remained positioned in later pseudotime regions, whereas DTAP_P, DTAP_T, DTAP_A, and DTAP_D populations were concentrated predominantly in early-to-intermediate pseudotime regions, although individual cells extended toward later pseudotime values. AT2 cells remained largely localized near the early pseudotime region. Thus, mOGG1 treatment modified the organization of alveolar epithelial states after chlorine exposure, with the distribution of DTAP populations remaining distinguishable from the later AT1-associated state.

Because the trajectory was rooted in control AT2 cells, the observed pseudotemporal ordering is consistent with progression from an AT2-like state through transitional DTAP-associated states toward an AT1-like state. However, the boxplots alone do not demonstrate a single linear path or prove that all DTAP populations directly differentiate into AT1 cells. Instead, they indicate condition-associated changes in the relative positioning and dispersion of alveolar epithelial states.

### Chlorine alters airway epithelial state organization

The airway trajectory showed a distinct organization from the alveolar lineage. In air-control lungs, basal cells were concentrated near the trajectory origin, consistent with their use as the root state (**Figure 5A**). Deuterosomal cells occupied an early-to-intermediate position, while secretory and ciliated cells were distributed at later pseudotime values. Secretory cells showed a relatively broad distribution with a subset of late-pseudotime cells, whereas ciliated cells occupied a more intermediate-to-late region.

**Figure 5.**
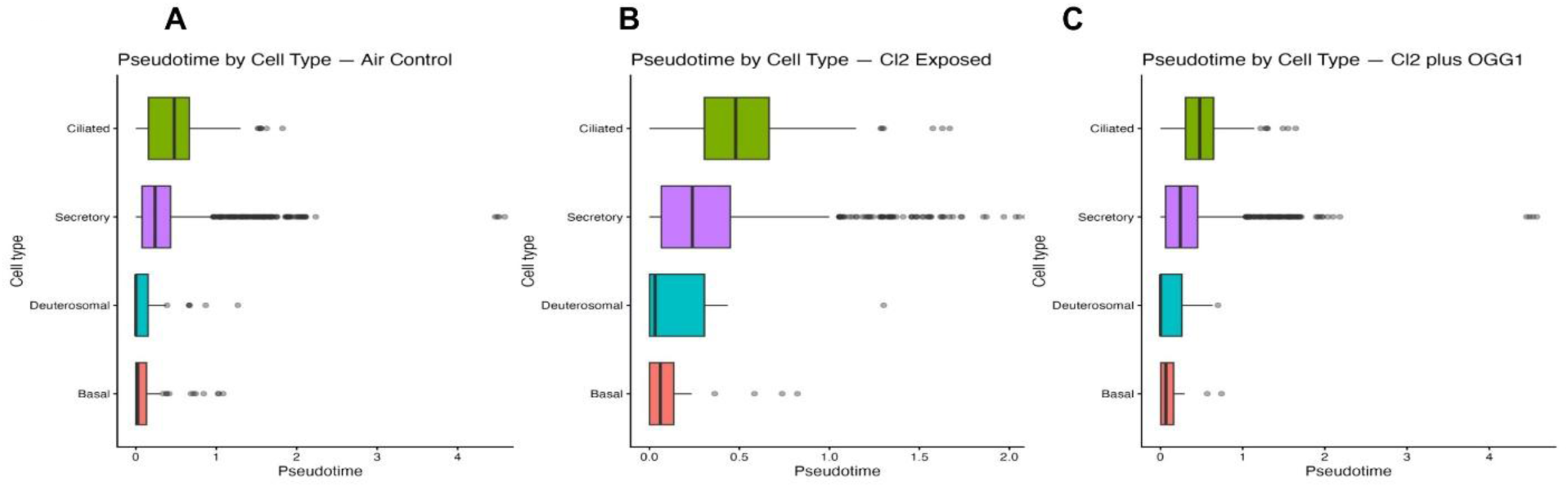
Pseudotemporal distribution airway epithelial states following chlorine exposure and mOGG1 treatment. Pseudotemporal analysis was performed for the airway epithelial lineages to evaluate injury-associated state transitions and potential movement toward repair-associated phenotypes. Pseudotime was inferred independently within each lineage. Boxplots show the distribution of pseudotime values for each annotated cell population, with individual points representing cells. **(A-C)** Airway epithelial trajectory analysis. The airway lineage included basal, secretory/club, ciliated, and deuterosomal cells. The trajectory was rooted in basal cells from air-control lungs. **(A)** Air-control lungs; **(B)** Cl₂-exposed lungs; and **(C)** Cl₂ plus mOGG1-treated lungs. In air-control lungs, basal cells occupied the earliest pseudotime region, followed by deuterosomal, secretory, and ciliated populations. Chlorine exposure altered the pseudotime distributions of secretory and ciliated cells, with increased dispersion and late-pseudotime outliers. Following mOGG1 treatment, the airway epithelial populations displayed a modified pseudotemporal distribution, with ciliated and secretory cells remaining the most advanced airway states. These analyses support condition-associated remodeling of alveolar and airway epithelial state progression after chlorine injury.

Chlorine exposure altered the distribution of airway epithelial states (**Figure 5B**). Ciliated cells showed a broader and more advanced pseudotime distribution than basal cells, while secretory cells displayed marked dispersion with numerous late-pseudotime outliers. Deuterosomal cells also showed a broader distribution than in air controls. These changes suggest that chlorine exposure disrupts the normal organization of the basal-to-differentiated airway epithelial continuum and induces heterogeneity within secretory and ciliated epithelial states.

Following mOGG1 treatment, the airway epithelial populations showed a modified pseudotemporal pattern (**Figure 5C**). Ciliated and secretory cells remained the most advanced airway populations relative to basal cells, while deuterosomal and basal cells remained concentrated in earlier pseudotime regions. The reduction in the apparent spread of some airway populations, together with persistence of late-pseudotime secretory cells, suggests partial but incomplete remodeling of the chlorine-associated airway response.

Together, the two lineage-restricted analyses demonstrate that chlorine exposure affects both alveolar and airway epithelial state organization, but in lineage-specific ways. The alveolar response was characterized by broadening of AT1 and DTAP-associated pseudotime distributions, whereas the airway response involved altered dispersion of secretory, ciliated, and deuterosomal states. mOGG1 modified these distributions in both lineages but did not completely restore them to the air-control pattern at 24 h. These findings support a model in which mOGG1 influences the organization of injury-associated epithelial states and may initiate a more controlled transition toward repair-associated phenotypes.

Because pseudotime is an inferred transcriptional ordering, these findings should not be interpreted as direct evidence of lineage conversion. Definitive confirmation of epithelial lineage progression and repair would require time-course analysis, lineage tracing, or complementary functional validation.

### mOGG1 redistributes chlorine-induced IL-1 signaling across immune and epithelial populations

Cell-cell communication analysis predicted a sparse IL-1 signaling network in air-control lungs (**Figure 6**). In air control group, neutrophils were identified as the principal IL-1 originators, with signaling directed mainly toward secretory epithelial cells and alveolar macrophages (AMs) (**Figure 6A**). Consistent with this pattern, network-centrality analysis identified neutrophils and AMs as the principal influencers and secretory cells as the dominant receivers. Other epithelial and myeloid populations contributed minimally to the basal Il-1 network (**Figure 6B**).

**Figure 6.**
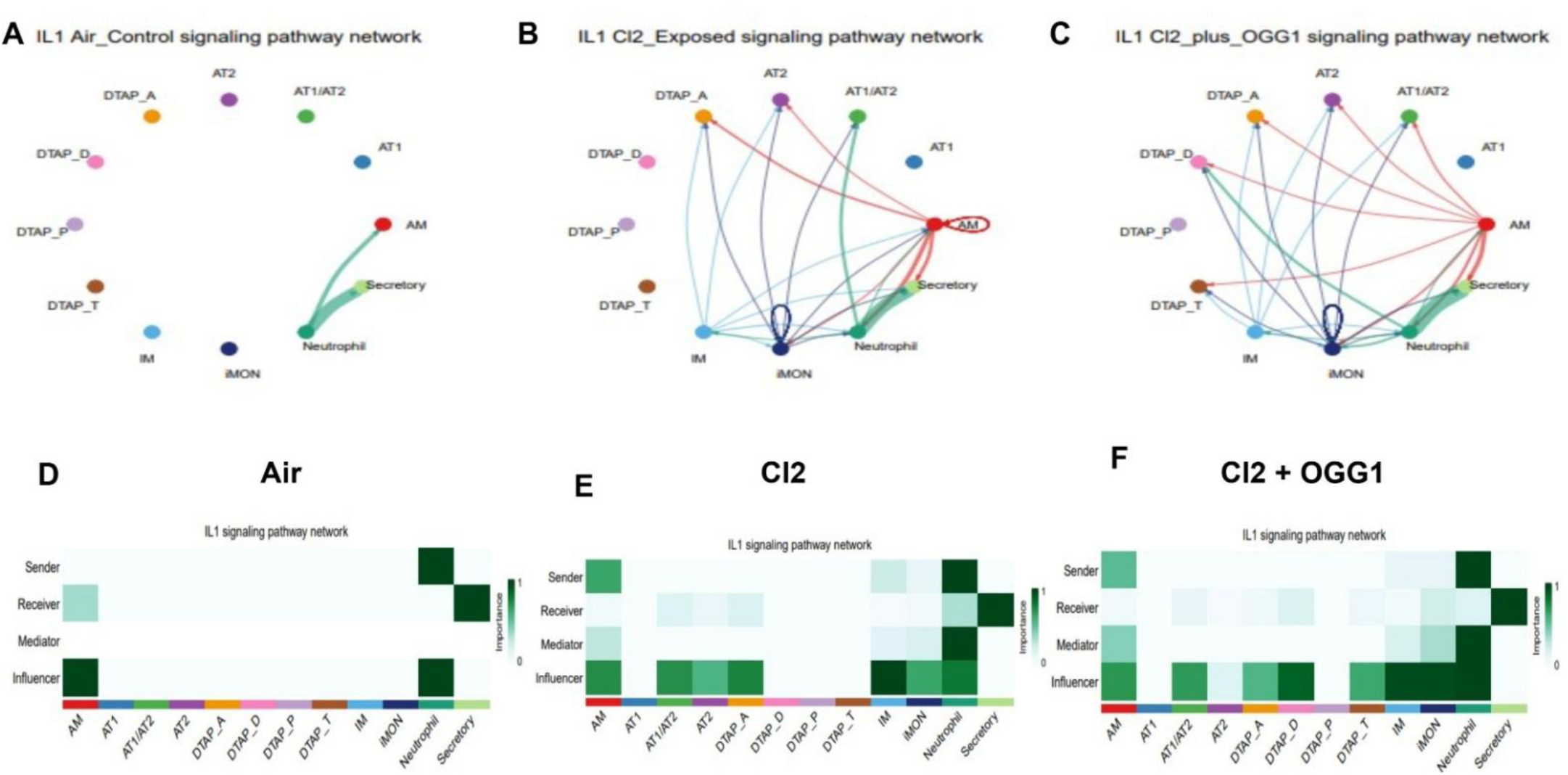
mOGG1 alters the IL-1 intercellular signaling network after chlorine exposure. Cell type-specific IL-1 pathway communication in lungs from mice exposed to room air **(A, B)**, **Cl₂ (C, D**), or Cl₂ +mOGG1 (**E, F**). Circle plots show the inferred IL-1 signaling network for each experimental condition (**A, C, E**). Each node represents an annotated lung cell population. Directed edges represent predicted ligand-receptor communication from a sending population to a receiving population; arrowheads indicate signaling direction, edge colors correspond to the sending population, and edge thickness represents the relative inferred communication strength. Self-directed edges indicate predicted autocrine signaling. The air-control network was comparatively sparse and was dominated by communication among alveolar macrophages, neutrophils, and secretory epithelial cells. Cl₂ exposure produced a broader IL-1 communication network involving myeloid, epithelial, and damage-associated transitional cell populations. The Cl₂ + mOGG1 condition retained extensive IL-1 communication but displayed a reconfigured pattern of predicted interactions among alveolar macrophages, inflammatory monocytes, interstitial macrophages, neutrophils, and epithelial populations. (**B, D, F**) Heatmaps summarize the relative network-centrality roles of each cell population in the corresponding condition. Rows indicate the inferred role of each population as a sender, receiver, mediator, or influencer. Columns represent the annotated cell populations. Darker green indicates greater normalized importance for the indicated signaling role, on a scale from 0 to 1; white or pale shading indicates little or no inferred contribution. The colored strip beneath each heatmap identifies the cell-population colors used in the corresponding circle plot. These analyses reflect transcript-based predictions of ligand-receptor communication and do not directly measure IL-1 protein production, secretion, or pathway activation.

Cl₂ exposure substantially broadened the predicted IL-1 network, engaging AMs, interstitial macrophages, inflammatory monocytes, neutrophils, and several other alveolar epithelial populations (**Figure 6C, D**). Neutrophils remained the dominant sender and mediator, whereas AMs acquired a prominent outgoing signaling role. Secretory epithelial cells remained the strongest receivers, with additional predicted incoming signals involving neutrophils and AT1/AT2, AT2, and DTAP_A cells. Interstitial macrophages, inflammatory monocytes, AMs, neutrophils, and multiple epithelial populations also displayed substantial influencer centrality. Thus, Cl₂ exposure transforms IL-1 communication from a limited baseline circuit into an extensive immune-epithelial communication network.

At 24 h after Cl₂ exposure, mOGG1 treatment did not eliminate the predicted IL-1 network but altered the distribution of signaling roles among participating populations (**Figure 6E, F**). Neutrophils remained the dominant sender and mediator, AMs continued to contribute to outgoing signals, and secretory epithelial cells remained the principal receivers. Interstitial macrophages and inflammatory monocytes also contributed as mediators or influencers. Notably, damage-associated transitional epithelial populations became prominent within the treated network: DTAP_D and DTAP_T cells exhibited strong influencer centrality, while DTAP_A, AT1/AT2, and AT2 cells remained integrated into the immune-epithelial signaling architecture. In contrast, DTAP_P cells showed little involvement under all conditions.

Together, these analyses indicate that mOGG1 did not globally suppressed the IL-1 communication at 24 h after Cl₂ exposure. Instead, treatment reorganized the predicted network while preserving neutrophils and AMs as major sources and secretory epithelial cells as a principal target. The increased centrality of selected damage-associated transitional population suggests that mOGG1 modifies how injured alveolar epithelial states participate in inflammatory communication. Because these interactions were inferred from ligand-receptor expression, they represent predicted signaling relationships rather than direct measurements of IL-1 secretion or pathway activation.

## Discussion

Alveolar repair requires coordinated interactions among AT2, AT1, immune, and stromal cells. After injury, AT2 cells can proliferate and differentiate toward an AT1 phenotype through transitional epithelial states expressing stress, remodeling, and alveolar genes, including *Krt8, Krt18, Krt19,* and *Cldn4*^22–28^ .Although these transitional states may support regeneration, their persistence has been linked to impaired epithelial differentiation, fibrosis, and chronic remodeling ^24–26,28^ . Oxidative mtDNA damage may contribute to these processes by activating mitochondrial dysfunction, cellular stress pathways, inflammatory signaling, and mitochondrial damage-associated molecular pattern responses ^8,21,29^ .

We previously showed that oxidative mtDNA damage contributes to chlorine-induced lung injury and that mitochondria-targeted OGG1 (mOGG1) improves mitochondrial function, epithelial ion transport, alveolar fluid clearance, and longer-term lung physiology after chlorine exposure^16^ . Here, single-nucleus RNA sequencing provided a cellular context for these effects. Twenty-four hours after chlorine exposure, mature AT2 representation was reduced, AT1/AT2 intermediate populations expanded, and distinct stress-responsive, *Krt8*-associated transitional, proliferative, and AT1-associated states emerged. These findings suggest that chlorine disrupts not only epithelial function, but also transcriptional programs involved in alveolar repair. This interpretation is consistent with prior studies showing that chlorine exposure causes oxidant injury, epithelial ion-channel dysfunction, impaired alveolar fluid clearance, and pulmonary edema ^30–33^.

Subclustering identified four differential transcriptomic adaptation states, DTAP_D, DTAP_T, DTAP_P, and DTAP_A, reflecting heterogeneous epithelial responses rather than a single linear repair pathway. Stress-related programs were prominent, while DTAP_T showed a *Krt8*-associated transitional profile, DTAP_P a proliferative profile, and DTAP_A a more AT1-associated profile. Similar *Krt8*⁺ transitional or damage-associated epithelial states have been described after lung injury and in fibrotic lung disease ^24–28^ . Pseudotime analysis was consistent with an injury repair continuum extending from AT2-like cells through stress-responsive and transitional states toward proliferation or AT1-associated differentiation. However, pseudotime reflects transcriptional similarity rather than elapsed time or direct ancestry. The inferred DTAP_D→DTAP_T→DTAP_P and/or DTAP_A organization should therefore be considered a working model, not proof of lineage progression ^34–36^.

Chlorine also altered airway epithelial populations and markedly increased the predicted complexity of IL-1-mediated communication among neutrophils, monocytes, secretory cells, AT2 cells, and transitional epithelial populations. IL-1 signaling has been implicated in epithelial injury responses, inflammatory amplification, leukocyte recruitment, and regulation of damage-associated alveolar progenitor states ^24,37,38^ This pattern is biologically plausible because IL-1 signaling promotes chemokine production, leukocyte recruitment, and epithelial plasticity, but persistent IL-1β signaling can impede terminal epithelial differentiation. Expression of *Nfkbia, Fos, Jun, Junb, Atf3,* and *Egr1* was also compatible with NF-κB and MAPK/AP-1 responses^37,38^ , although transcript abundance alone does not establish pathway activation.

Expression of *Ccl2, Cxcl1,* and *Cxcl2* was consistent with potential recruitment of monocytes and neutrophils through CCL2-CCR2 and CXCL1/CXCL2-CXCR2 pathways^39,40^. However, ligand-receptor analysis is computational and cannot establish signaling chronology, protein activation, or functional signaling without experimental validation ^41,42^ .

mOGG1 did not globally suppress predicted inflammatory communication. Instead, under the specific conditions of this study: a single chlorine exposure, one intranasal mOGG1 dose 1 h after exposure, and analysis at 24 h, mOGG1 partially reduced AT1/AT2 intermediate-cell accumulation, constrained selected proliferative programs, and redistributed epithelial-immune interactions, including IL-1 signaling. This pattern suggests an actively reorganized early injury-repair response rather than complete resolution at 24 h.

This study has several limitations. The single 24h time point cannot determine whether DTAP states resolve, persist, or predict the chronic physiological abnormalities observed after chlorine exposure. Pseudotime analysis cannot establish lineage relationships, and ligand-receptor analyses are computational predictions requiring experimental validation. Changes in cell representation in single-nucleus datasets also may reflect cell loss, altered transcriptional identity, or differences in nuclear recovery rather than absolute changes in abundance. Finally, although both sexes were represented equally, the study was not powered to detect sex-specific effects.

In conclusion, acute chlorine exposure was associated with altered alveolar and airway epithelial organization, expansion of AT1/AT2 transitional states, and increased predicted immune-epithelial communication. Under the specific conditions examined in this study with a single chlorine exposure, one intranasal dose of mOGG1 administered 1 h later, and analysis at 24 h, mOGG1 partially reduced these epithelial changes and altered predicted IL-1 signaling without restoring the lung to the air-control state. These data define early cellular and transcriptional responses associated with chlorine exposure and mOGG1 treatment but do not establish that oxidative mtDNA damage directly causes altered epithelial plasticity or inflammatory communication, nor that mOGG1 acts through these pathways to promote repair. Longitudinal, lineage-tracing, and functional studies are needed to determine the fate of DTAP populations, validate the predicted signaling interactions, and establish their mechanistic relationships to mtDNA damage and long-term lung outcomes.

## Acknowledgement

We are grateful to Dr. Mark Gillespie (University of South Alabama) for generously providing the mitochondria-targeted OGG1 (mito-OGG1) recombinant protein used in this study.

## Author contributions

S.M. and S.D. conceived and designed the research. S.D. performed the experiments. D.K.C. and S.D. helped with data analysis. S.D., D.K.C., and S.M. interpreted results. S.D. drafted the manuscript; S.M. and D.C. edited the manuscript. All authors discussed the manuscript at each stage and approved of the final version.

## Competing interests

None of the authors have a competing interest.

## Materials & Correspondence

Correspondence and material requests can be addressed to Dr. Sadis Matalon.

## Sources of Funding

These studies were supported by 3R01HL031197-29S1, 1R61HL179700-01A1 and 5R01HL031197-31 to Dr. Sadis Matalon.

## Notes

### Competing Interest Statement

The authors have declared no competing interest.

